# On the origin of nonequivalent states: how we can talk about preprints

**DOI:** 10.1101/092817

**Authors:** Cameron Neylon, Damian Pattinson, Geoffrey Bilder, Jennifer Lin

**Affiliations:** Centre for Culture and Technology, Curtin University; Research Square; Crossref

## Abstract

Increasingly, preprints are at the center of conversations across the research ecosystem. But disagreements remain about the role they play. Do they “count” for research assessment? Is it ok to post preprints in more than one place? In this paper, we argue that these discussions often conflate two separate issues, the history of the manuscript and the status granted it by different communities. In this paper, we propose a new model that distinguishes the characteristics of the object, its “state”, from the subjective “standing” granted to it by different communities. This provides a way to discuss the difference in practices between communities, which will deliver more productive conversations and facilitate negotiation on how to collectively improve the process of scholarly communications not only for preprints but other forms of scholarly contributions.

## I. Introduction

Two scientists, Jimmy Maxwell and Chuck Darwin, meet at a conference and realise that, although one is a physicist and the other a naturalist, they have common research interests, and agree to collaborate. Their work develops quickly and a theory emerges that could revolutionise both their disciplines.

They write up their work and, egged on by the physicist, decide to post to a preprint server before formally submitting to their target journal, *The Science of Nature.* The preprint causes a sensation! It receives attention, generates heated discussion, and citations ensue from their colleagues in both disciplines. The journal submission, however, faces a rockier path, getting held up by Reviewer #3 through four rounds of revision over a sticky issue involving the techniques for measuring the forces of barnacle-rock attraction.

During the publication delay, offers start pouring into young Maxwell’s inbox from universities and companies wishing to recruit the young physicist to develop his findings. He takes a plum job and goes on to change the course of physics forever. Chuck, on the other hand, finds offers harder to come by. His applications for grants to fund a trip to far-flung islands fail because his CV lacks the high impact articles required to make him stand out from the other candidates. In despair he quits the bench and opens a pet shop. Some decades later the two researchers are recognized by the award of the prestigious Prize of Nobility. Maxwell’s place in the firmament is assured, while Darwin returns to his pet shop, now specialising in finches, where something about their beaks bothers him until the day he dies.

We offer this cheeky story to illustrate one main point: different communities grant the same object different degrees of importance. We can complicate the story by revealing that both researchers were scooped between posting the preprint and article publication. Or funding panels, one in bioscience and the other in physics, assess their applications, counting the outputs as scholarly contributions in different ways. But we are illustrating the same point. There is no universal standard of when an output is considered as part of the formal scholarly record. Rather, it is determined by particular groups in particular contexts.

### The problem of defining “a preprint”

The pace of technological change over the past two decades has far outstripped the language we use to describe the objects and processes we use to communicate. This disconnect between language and technology is at the root of the current debate around preprints. The very word “preprint” is an odd combination of retronym and synecdoche, increasingly unlikely to ever be a precursor to anything that appears in physical print. At the same time, the root of the word remains “print,” which takes one small part of the publishing process to stand in for the entire process. A preprint is different from a working paper, yet both are entirely different to an academic blog post. Additionally, all these appear in designated online repositories as digital documents that are recognizably structured as scholarly objects. Some preprints are shared with the future intent of formal publication in a journal or monograph. But not all. And the term is used to mean a host of different things, and as such, remains referentially opaque.^1^

Wikipedia is a good source for identifying common usage. At the time of writing it defines a preprint as “a draft of a scientific paper that has not yet been published in a peer-reviewed scientific journal.”[1] This definition encompasses everything from early, private drafts of a paper that the author(s) have never shared with anyone all the way to drafts of accepted manuscripts that have yet to go through a publisher’s production process. Interpreted liberally, the Wikipedia page itself might even be included.[2] The definition also conflates science and scholarship in a way which is both common and unhelpful. For many readers it would exclude work from the social sciences and humanities, as well as book chapters and other drafts destined for venues beyond “a peer-reviewed scientific journal”.

Other organizations have constructed their own meanings and terms to fit the agenda of their constituencies. SHERPA, a UK organisation dedicated to studying scholarly communication, has a more precise definition for preprints: “the version of the paper before peer review”.[3] They then define versions between acceptance and publication as “post-prints.” NISO (National Information Standards Organisation) doesn’t formally define the word “preprint” in its Journal Article Version (JAV) standard[4], preferring instead to further delineate where “significant value-added state changes” occur. They break down the broad Wikipedia definition into four distinct stages including “author’s original”, “submitted manuscript under review”, “accepted manuscript” and any numbers of “proofs” that may emerge between acceptance and the published “version of record”, a term which suffers under the dual burden of being both essentially undefinable and highly politicised.

As a further complication, the shifting roles of different players in the ecosystem have also contributed to this confusion. To “publish” a work can mean three entirely different things: the labour of preparing a work for its dissemination, to communicate or make public a work, or in the narrow sense we use in the academy, to make available through designated channels after specified social and technical processes. “Preprint” is positioned and often defined in relation to “publish” in a way that adds to the ambiguity of both terms.

In the past there was a clear distinction between services that hosted preprints and “publishers” who carried out the formal process of “publication” as defined by scholarly communities. A preprint could therefore be identified by its presence on a platform that was not that of a “publisher”. But today, publishers are starting to provide repositories to host preprints (PeerJ, Elsevier/SSRN, and the American Chemical Society). To add to the confusion, new forms of journals that run quite traditional quality assurance and review processes are being developed, which use preprint servers as the storage host for their articles. *Discrete Analysis*[5] and *The Open Journal*[6] both use ArXiv to store the PDF versions of accepted papers. A definition that depends on the historical role of any given player will fail if that role changes. Attempts to define the term “preprint” in this way push the confusion onto other, equally poorly defined, terms. Saying a preprint “is not published” or “is not in a journal” shifts the ambiguity to the question of what “published” means or what counts as a “journal.”

The lack of clear definitions is a problem when discussing and negotiating important changes to research communication. Researchers today can share results earlier, in new forms, and to new communities. But the newness of such technologies means that we have not yet come up with terminology to clearly discuss available choices. Some researchers simply see a preprint as an early notification or preview of a “formal” publication. For others it is a complete finding and a clear claim of priority in the scholarly literature. These differences are most often due to differences in disciplinary cultures. And, as in our story, the confusion is even greater with work that crosses disciplinary boundaries.

This is fundamentally an issue of what “counts”, and what counts will clearly depend on the community that is doing the counting. This is the central social and political issue that disagreements on the status of preprints circle around. Our argument is that in these discussions we unhelpfully conflate this issue of the “standing” of an object – whether it counts – within a community with the related but separate issue of the characteristics of the object itself. What processes has it been through? How has it been validated or checked? By separating questions of the “state” of the object from the “standing” that different communities give it we can have much more productive discussions and negotiations about what will count, and why.

## II. The State-Standing Model

### State vs. Standing

While “preprints” is a referentially opaque term that make little sense in the context of an online communications environment, it is unlikely we will persuade anyone to abandon the term. Instead, we seek to tease out two of its attributes often elided when discussing any scholarly communication: “state” and “standing.”

##### Attributes of a Research Object

- *State* - the external, objectively determinable characteristics
- *Standing* - the position, staius, or reputation

The “**state**” of a research object is comprised of the external, objectively determinable characteristics of the object. This includes records of claims made about the object, metadata, statements of validation processes the object has undergone, etc. A manuscript undergoes a wide array of state changes as multiple players interact with it in the process of submission and publication: technical checks and validation, editorial assessment, assignment of editor and reviewers, referee review, editorial decision, typesetting, author approval and corrections, publication accept, content registration/metadata depositing, front matter editorial posting, publication commentary facilitation, retraction/correction processes, publication event tracking, etc. This includes explicitly modelled metadata elements within strong schema (such as “indexed in PubMed”) as well as unstructured and vague terms. It also includes a description of groups that have access, including “the public”.[7] With “state,” there *can be* an explicit record made even if it is not exposed. It may be hidden within publisher systems. Or it may be private information unethical to share. The record might be in third party systems such as Pubmed Central or ORCID. Or some elements may be badly recorded or lost, and thus inaccessible.

If a manuscript changes state, it may also also undergo changes in value or intellectual status. The “**standing**” of a research object is the position, status, or reputation of an object. It is a consequence of its history and state. There are various forms of standing recognised by different groups, for example: “has been validated by (a traditional) peer review process”, “establishes priority of claim”, “is appropriate for inclusion in this assessment process,” “is considered appropriate for discussion and thus citable”, etc. These are judgments about the recognition or value of the output. Standing is conferred by a group, not an individual, and is therefore distinct from any individual’s opinion of the work.^2^ It is also conferred not directly to individual objects but to classes of objects that share attributes of state.

### Nonequivalent states, nonequivalent changes

With a conceptual barrier between state and standing in place, we can investigate their relationship as the scholarly output changes over time. A state change may lead to a change in standing, but not necessarily and not in all cases. A change in standing, however, only occurs as a consequence of a state change triggered by some external shift that has led to a reconsideration of value.

Standing is independently conferred by each group for whom the research output has meaning. While *similar* forms of standing between groups might arise, they cannot be identical as such. What matters most in this model is the possibility that a particular community may confer a different form of standing than another on the same type of research object (i.e., with the same state).

Figure 1 shows that research changes state, but to different effects in changes of standing across physics and life science communities. Both may consider research validated and part of the formal record at similar stages of the publication process. But there are also key differences. When a preprint is posted by a physicist, s/he has established the priority of claim in that community. And it is considered worth of citation. For the life sciences community, however, claim priority is generally established when a manuscript is submitted to a journal. It is only appropriate to cite the article even later, when the manuscript is made available online (Advanced Online Publication or online publication).

**Figure 1:**
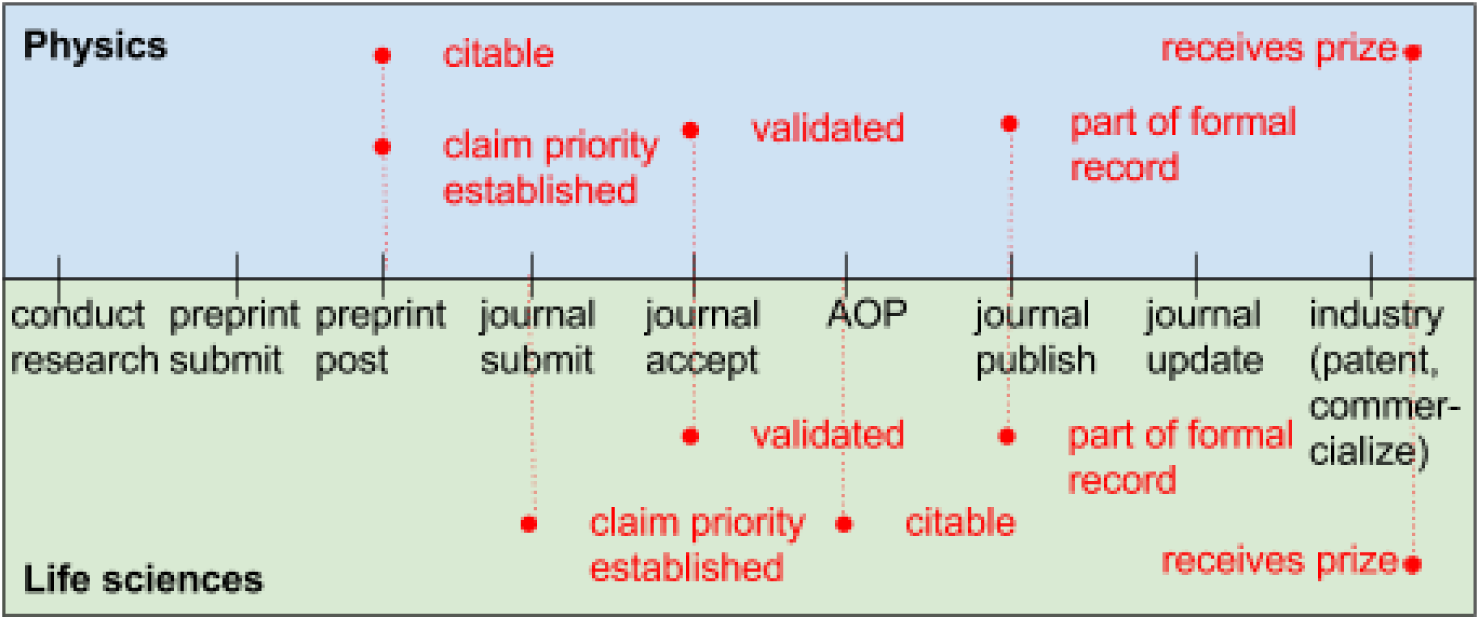
*Differences in standing (red) between Physics and Life science communities*.

The conditions which prevail in the conduct of research are naturally tied to the type of research itself. As these vary widely, so would the influence they have on the communication culture of the group and how they confer status. That certain fields in physics share equipment, work in very large groups, etc. has been often mentioned as a contributor to their predilection for preprints. On the other end of the publication event, research may expand its reach and utility beyond the academy. This introduces other possible entities that begin to serve as a conferrer of status (ex: university office of technology transfer), and it will vary by field and discipline depending on the opportunities possible. Both Maxwell and Darwin are awarded for their work in acknowledgment of their contributions. But given that the research was taken up by the physics community earlier, it would not be surprising to see time differences in the subsequent accolades offered to each by their respective disciplines.

### Applying the model to preprints

Prior to the development of the web, some segments of both Economics and High Energy Physics communities shared a similar practice, the circulation by mail of manuscripts to a select community, before submission for formal peer review at a journal. As the web developed, both communities made use of online repositories to make this sharing process more efficient and effective. Paul Ginsparg initially created ArXiv as an email platform, but then migrated it onto a web-based platform in the early 1990s. In 1994, two economists created the Social Sciences Research Network (SSRN), a platform that shared many traits with ArXiv. In both cases researchers submit digital manuscripts, which undergo a light check prior to being made publicly available on the platform. These manuscripts have not been subjected to any formal version of review by expert peers. Furthermore, there is a common expectation in both repositories that most manuscripts will go on to be formally published as journal articles or book chapters. That is, the *state* of objects in both ArXiv and SSRN is very similar.

Nonetheless the *standing* of these objects for these two communities are quite different. For the High Energy Physics community (and others in theoretical physics), posting to ArXiv establishes the priority of claims and discoveries. In many ways ArXiv preprints are seen as equivalent to formally published articles, and many physicists will preferentially read articles at ArXiv rather than find copies in journals. Indeed, for those disciplines where use of ArXiv is common, the formal publication is the point at which citations to the manuscript start to drop off.[8] The question of why physicists continue to publish in journals at all is a separate one and beyond the scope of this paper. However our model can help: clearly the community, or communities that matter, do grant some standing to journal articles which is both different to that granted to preprints and important in some way. The question of what that standing is and why it continues to matter is separated in our model from the equivalence of state that journal articles in physics share with those in other disciplines. As Maxwell and Darwin found in our story physics and biosciences are different in important ways, even when their publication processes are very similar.

By contrast, working papers on SSRN are seen much more as works in progress. They are frequently posted well before submission to a journal, unlike ArXiv where posting is frequently done at the same time as submission. Observers from outside these communities, including those interested in adopting physics posting practices for the biosciences, often make the mistake of seeing two similar repositories with similar requirements and assume that SSRN working papers and ArXiv preprints can be equated. The differences are not obvious from an examination of *state*, but are situated in differences in standing. Working papers and preprints have a different *standing*, and serve quite different functions for their cognate communities, despite being quite similar in form. Separating the two concerns allows us to be much clearer about what is similar and what is different between the two cases.

## III. Further applications in the publishing life cycle

The uses of our model are not limited to preprints. It is a useful heuristic for isolating the questions that require answers from a community, from those that can be answered by auditing the process an object has been through. That is, it is helpful to separate the question of whether something has been done, from the question of whether any community cares.

We believe this separation of concerns will be valuable for discussions on a wide range of outputs including software, data and books. Indeed all types of research outputs go through processes of validation, dissemination and assessment, which are accorded differing degrees of importance by different communities. Discussions of the details of options for differing modes of open, signed, partially open, single blind, double, or even triple blind, peer review will benefit from separating the description of process (and testing whether the stated process has been followed) from the views of any given community of objects that have been through that process.

It may be the case that much of the confusion around newer forms of scholarly sharing, including efforts to make certain scholarly outputs “matter” as much as traditional narrative publications is due to this same confusion. New forms of output seek to co-opt the expression of forms of state, without putting in the required work that connects the social machinery of state-standing links. As a result they frequently fall into an “uncanny valley”, objects that look familiar but are wrong in some subtle way. The most obvious example of this are efforts to make new objects “citable”, i.e. making it technically feasible to reference in a traditional manner through provision of specific forms of metadata, most commonly via DOIs. To actually shift incentives this work needs to be linked to a social and political shift which changes a community’s view of what they should cite, i.e. what gives an object sufficient standing to make it “citation-worthy”.

A similar debate is that which rages between traditional publishers and advocates of a shift towards “publish-first, review-later” models of research communication. One one hand, advocates of change often remark on the seeming lack of improvement made to the *text* of an article through traditional peer review. For example Klein et. al. found that text content of ArXiv preprints only undergo minor changes between the initially submitted and finally published versions.[9] Of course this neglects state changes in the validation process that may be important but are not necessarily reflected in the character-stream of the article, such as ethical or statistical checks that were passed.

On the other hand, publishers have established practices that they consider important, captured in the JAV vocabulary.[4] JAV details a number of different stages (with different states), which a manuscript might go undergo. Many of these are invisible to authors. For instance, Author Original and Submitted Manuscript Under Review are identified as distinct states. An author would consider these to be the same document, but a publisher needs to record the manuscript’s transition into the peer review pipeline. At the same time, JAV ignores changes that are likely of concern to authors by failing to record them. For example it has no concept of the distinct revised versions of a manuscript submitted during review cycles.

Here, the issue is a difference in focus on what it is that matters, what kinds of standing are important. Changes in state that are important markers of shifts in standing for one group are ignored by the other and vice versa. Until the full set of state changes that are relevant to all stakeholders are transparently visible, discussions of standing are unlikely to be productive.

This illustrates a crucial point. Our model exposes the need for high quality metadata, that is well coupled to the record of processes that a work has experienced. If what is contained within the scholarly record is a question of standing, then the formal record of state is a critical part of supporting claims of research.

## V. Conclusion

To engage in productive discourse on new (and traditional) forms of scholarly sharing, we need to gain clarity on the objects themselves. We propose a model that explicitly separates the state of a work – the processes it has been through and the (objectively determinable) attributes it has collected through those processes – from the standing granted it by a specific community. It is not only a formal framework but a practical apparatus for navigating and negotiating the ongoing changes in scholarly communications. By distinguishing two attributes we can isolate aspects of objects that can be easily agreed on across communities, and those for which agreement may be difficult. These have clouded discussion of community practices, particularly those around the emerging interest in “preprints” in disciplines that have not previously engaged in the sharing of article manuscripts prior to formal publication.

How does our model help the young Darwin and Maxwell? At one level it doesn’t. Questions of standing will be inherently difficult to discuss across community boundaries. It cannot solve the underlying *social* challenge that different research communities simply value different things, and in particularly different parts of the overall life cycle of a research communication. But each community also benefits from a discussion that makes the currently implicit connection between state and standing explicit. It offers a way of talking about and analysing those differences, differentiating between changes to the object and changes to the perception of that object. To bring the culture of manuscript posting to the biosciences, Darwin would be better served by identifying the different goals that different people had as well as discuss the concerns that more traditional researchers have.

Publishers, including preprint repositories, can better serve their communities by making state changes much clearer, more explicit, and transparent. It is impossible for us to make progress in discussing standing when we cannot clearly define what the state is. We cannot discuss the difference in standing between a preprint, a journal editorial, and a research article without knowing what review or validation process each has gone through. A further consequence of our argument is that service providers need to pay much greater attention to recording state changes that matter for the communities that they serve, rather than only those that are for their internal use.

But it is for scholarly communities to grant standing as a result of the clear and formal record of social processes through which they validate scholarly work. By making explicit both the distinction between social processes and the record of attributes that results from them, and explicitly recognising the connection between state and standing, we surface the processes of scholarship more clearly, and re-centre the importance of communities deciding for themselves what classes of object deserve which granting of standing.

## Footnotes

1 It should be noted that this is not a new problem. For many years researchers have bundled everything that has not been formally “published” under the umbrella term, “grey literature”, creating headaches for every metaanalyst and systematic reviewer who has had to decide what “counts” as a meaningful academic contribution.

2 We use the general term “group” to refer to communities, institutions and other parties that confer standing. “Groups” therefore includes disciplinary communities, universities (and their departments), funders, but also potentially entities such as main stream media venues as well as specific publics.

